# Collagen content and cross-links scale with passive stiffness in dystrophic mouse muscles, but are not altered with administration of collagen cross-linking inhibitor beta-aminopropionitrile

**DOI:** 10.1101/2022.07.08.499292

**Authors:** Sarah E. Brashear, Ross P. Wohlgemuth, Lin-Ya Hu, Elias H. Jbeily, Blaine A. Christiansen, Lucas R. Smith

## Abstract

In Duchenne muscular dystrophy (DMD), a lack of functional dystrophin leads to myofiber instability and progressive muscle damage that results in fibrosis. While fibrosis is primarily characterized by an accumulation of extracellular matrix (ECM) components, there are changes in ECM architecture during fibrosis that relate more closely to functional muscle stiffness. One of these architectural changes in dystrophic muscle is collagen cross-linking, which has been shown to increase the passive muscle stiffness in models of fibrosis including the *mdx* mouse, a model of DMD. We tested whether the intraperitoneal injections of beta-aminopropionitrile (BAPN), an inhibitor of the cross-linking enzyme lysyl oxidase, would reduce collagen cross- linking and passive stiffness in young and adult *mdx* mice compared to saline-injected controls. We found no significant differences between BAPN treated and saline treated mice in collagen cross-linking and stiffness parameters. However, we observed that while collagen cross-linking and passive stiffness scaled positively in dystrophic muscles, collagen fiber alignment scaled with passive stiffness distinctly between muscles. We also observed that the dystrophic diaphragm showed the most dramatic fibrosis in terms of collagen content, cross-linking, and stiffness. Overall, we show that while BAPN was not effective at reducing collagen cross- linking, the positive association between collagen cross-linking and stiffness in dystrophic muscles still show cross-linking as a viable target for reducing passive muscle stiffness in DMD or other fibrotic muscle conditions.

**Key Points:**

- BAPN did not reduce collagen cross-linking or passive stiffness in any muscle
- Collagen cross-links scaled with passive stiffness in dystrophic muscles
- The *mdx* diaphragm showed the most dramatic fibrosis related to collagen content, cross- linking, and passive stiffness
- Collagen fiber alignment scales with passive stiffness differently between muscles

## INTRODUCTION

The mechanical function of skeletal muscle depends on the active contractile properties and the passive mechanics, both of which are compromised in muscle disease. Although there are several structures that contribute to the passive mechanics of skeletal muscle, one of the primary determinants is the extracellular matrix (ECM) (1–3). The ECM contributes to passive muscle stiffness largely through fibrillar collagen, specifically type I and type III (4). Collagen has high tensile stiffness which produces passive tension in unstimulated muscle under strain and transmits the active forces produced by muscle contraction (5, 6). The healthy ECM is critical to muscle function, however, in skeletal muscle diseases characterized by fibrosis such as Duchenne muscular dystrophy (DMD), there is a pathologic accumulation of ECM components which can cause excessive stiffness and often lead to joint contracture (7, 8). Muscle contractures and loss of range of motion develop frequently in muscular dystrophies as a consequence of high muscle passive stiffness (9, 10). Although contracture is a common symptom of fibrotic diseases, different muscles exhibit various degrees of dysfunction. The diaphragm is consistently the most severely affected by fibrosis, which contributes to respiratory failure and early mortality (11).

Although there are some surgical and physical remedies available to alleviate contractures of limb muscles, none of them reverse fibrosis and long-term efficacy is limited (12). Therefore, there is a need to elucidate the molecular structures responsible for excessive muscle stiffness in order to develop targeted therapies for contracture in neuromuscular disorders.

While blocking collagen expression in rodent models of DMD has been accomplished (13), there are off-target effects and no anti-fibrotic therapy is approved for neuromuscular conditions.

While the amount of collagen in muscle can contribute to its mechanical properties, other features of collagen architecture have been shown to be more critical in determining muscle stiffness. We have previously demonstrated that more nuanced qualities of the ECM, such as alignment, also hold strong predictive ability for both dynamic, velocity-dependent, and elastic, velocity-independent, passive muscle stiffness (14, 15). Collagen cross-linking is feature of ECM organization that is known to contribute to stiffness and has been shown to increase in DMD and aged muscle (16, 17). Collagen cross-linking inhibitors have been applied successfully in other fibrotic tissues (18–20), but have not been applied to skeletal muscle. Cross-links can be enzymatically formed through lysyl oxidase (LOX) and associated enzymes or through advanced glycation end-products (AGE) (18,21,22). The frequency of collagen cross-linking varies between different muscles as does the plasticity of cross-link frequency in fibrosis (16). Given their association with muscle passive stiffness and fibrosis, collagen cross-links are an attractive target for contracture therapy.

In this study we focused on inhibition of collagen cross-link formation as a therapeutic to reduce passive stiffness in skeletal muscle of *mdx* mice, a model for DMD. We used beta- aminopropionitrile (BAPN) as a naturally occurring inhibitor of the cross-linking enzyme LOX (23). BAPN has been used in other mouse studies and shown to be well-tolerated and effective at reducing collagen cross-links in tissues other than muscle (20). Importantly, LOX based collagen cross-linking is critical to bone integrity (24), but can be blocked with BAPN and is functionally impaired in dystrophic mice (25). The *mdx* mice on the DBA/2J genetic background produce more severe fibrosis, thus more similar to the human condition than the *mdx* mice on the C57BL/6 background (26). As fibrosis in the *mdx* DBA/2J strain is progressive (27), we utilized cohorts of young and adult mice to assess the preventative and reversal effects of BAPN treatment, respectively. While BAPN treatment was unsuccessful in inhibiting collagen cross- links in muscle, this study strengthens the muscle dependent association of ECM architecture with muscle stiffness.

## MATERIALS AND METHODS

### Ethical Approval

All animal experiments were approved by the University of California, Davis Institutional Animal Care and Use Committee.

### Animal Handling and BAPN administration

DBA/2J (wildtype) and D2.B10-Dmd*mdx*/J (*mdx*) mice were housed and bred in the UC Davis Teaching and Research Animal Care Services (TRACS) facility. They were grouped and housed on a 12:12 light-dark cycle, and given ad libitum access to food and water. The mice (adult wildtype male N=4, adult *mdx* male N=7, young wildtype male N=6, young *mdx* male N=5, adult wildtype female N=5, adult *mdx* female N=6, young wildtype female N=6, young *mdx* female N=6) were between 16-19 weeks (adult) and 4-5 weeks (young) of age when BAPN or PBS treatment was initiated. 100 mg/kg/day of BAPN was administered through daily intraperitoneal injections for 4 weeks prior to sacrifice. Body mass was measured weekly.

### Muscle Isolation

Mice were anesthetized using 2.5% Isoflurane gas in 1 L/min of oxygen. After isolation of lower limb muscles, euthanasia by cervical dislocation was administered while mice were under anesthesia. Soleus, extensor digitorum longus (EDL), and diaphragm muscles were collected for isolated muscle contractile and passive mechanical measurements. Muscles were stored in Ringer’s solution (Sodium Chloride, Potassium Chloride, Calcium Chloride dihydrate, Potassium Phosphate Monobasic, Magnesium Sulfate, 4-(2-Hydroxyethyl)piperazine-1-ethanesulfonic acid, Glucose) supplied with oxygen directly following collection and during mechanical testing. After passive and active mechanical testing, muscles from the left limb were flash frozen in liquid nitrogen and stored at -80°C while muscles from the right limb were pinned at the estimated optimum muscle length (L_o_) and fixed in 4% paraformaldehyde (PFA) for one day before storage in 4°C PBS.

### Passive Muscle Mechanics

The passive mechanical properties were measured as previously described (14). Briefly, following isolation of EDL and soleus, 7-0 sutures were tied at the muscle-tendon junctions. A small strip of diaphragm was separated with knots tied at the central tendon and the ribs. The suture loops were secured to the 300C-LR-Dual-Mode motor arm and force transducer (Aurora Scientific) in a well of 20°C Ringer’s solution with bubbling oxygen (28, 29). The L_o_ was ascertained through a series of twitches as the muscle was gradually stretched using the 701C stimulator (Aurora Scientific), with the length set at the maximum twitch amplitude (30). The L_o_ length was measured using a caliper between the site where the sutures were tied (the proximal and distal muscle-tendon junctions). The muscle length (L_f_), mass (m), ratio of fiber length to L_o_ (L_f_/L_o_), and density (ρ=1.06 g/cm^3^) were utilized to find the physiological cross-sectional area (PCSA) (31): PCSA=m/(L_o_·(L_f_/L_o_)·ρ).

The strains that took place in the passive mechanical protocol were derived from the L_o_. Before each strain the muscle was conditioned by periodic lengthening to the given strain at 1 Hz for 5 seconds (s). After conditioning, the muscle was stretched 2.5% at 1 L_o_/s and kept at the length for 120 s. Conditioning, strain, and relaxation was repeated at 5, 7.5, 10, and 12.5% strain. The elastic stiffness is the slope of the quadratic fit of elastic stress, the stress after 120 s of relaxation, at 10% strain (15). Muscles that failed before the 10% strain were eliminated from the sample. This technique is similar to other prior studies (15,32–34).

### Active Muscle Mechanics

Active mechanical protocols were performed on EDL and soleus muscles following passive mechanical measurement. The active mechanical protocol consisted of measuring the maximum isometric twitch force, allowing a 30 s rest, and measuring maximum isometric tetanus. This process was replicated two more times with 5 minutes between each bout of twitch and tetanus. The highest twitch and tetanus of the three trials was used in analysis.

### Hydroxyproline Assay

EDL, soleus, and diaphragm muscles from the left limb were measured for collagen content and cross-linking using the hydroxyproline and collagen solubility assay similar to other studies (15,35,36) and as described previously (14). To begin the hydroxyproline assay, muscles were removed from -80°C storage and powdered with mortar and pestle. Any remaining tendon tissue was removed from the muscle during powdering. The muscle powder was weighed, washed in 1 ml of PBS, and stirred for 30 minutes at 4°C. After stirring, the powder and solution were centrifuged at 16,000g for 30 minutes at 4°C. Following centrifugation, collagen that was not cross-linked was digested in a 1:10 (mass:volume) solution of 0.5M acetic acid containing 1 mg/ml pepsin, and was stirred overnight at 4°C. The following day, the sample was centrifuged at 16,000 g for 30 minutes at 4°C and the supernatant and pellet were separated. The supernatant contained the pepsin-soluble fraction (PSF) while the pellet contained the pepsin-insoluble fraction (PIF). Both the PSF and PIF were hydrolyzed in 0.5 ml of 6M HCl at 105°C overnight. 10 μL of each sample was combined with 150 μL isopropanol and 75 μL of 1.4% chloramine-T (ThermoFisher) in acetate citrate buffer and oxidized for 10 min at room temperature. The samples were combined with 1 ml of 3:13 dilution of Erlich’s reagent [1.5 g of 4- (dimethylamino) benzaldehyde (ThermoFisher); 5 ml ethanol; 337 μL sulfuric acid] to isopropanol and kept for 30 minutes at 58°C. Each assay included a standard curve (0-1,000 μM trans-4-hydroxy-_L_-proline; Fisher). Data were reported as μg hydroxyproline per mg powdered tissue wet mass.

### Histological Collagen Architecture

Picrosirius red staining was done similarly to other studies (14,15,37,38). Before sectioning, fixed muscles were embedded in 4% agarose. 200 μm thick longitudinal sections were sliced using Leica VT1000S. Sections were washed, dried for 60 min, and stained for 60 min in 0.1% (weight/volume) Direct Red 80 (Fisher) dissolved in saturated aqueous picric acid (Fisher).

Sections were washed twice for 60 s in 0.5% acetic acid, then dehydrated with three 60 s washes of 100% ethanol. The sections were then cleared using CitriSolv (Fisher Scientific) for 3 min, and blotted with Permount (Fisher Scientific).

Full longitudinal sections were imaged via a 20X objective with brightfield illumination on a Leica DMi8 microscope and DFC9000GTC camera. Linearly polarized light imaging required a rotating polarizer in the beam path before and after the sample. A sequence of ten tiling scans were imaged at angles from 0 to 90° in increments of 10°. ECM architecture was quantified using a custom MATLAB script providing MicroECM alignment and MacroECM deviation parameters as described previously (14) in conjunction with the polarized light images.

### Second Harmonic Generation Microscopy Collagen Architecture

Second Harmonic Generation (SHG) microscopy samples were collected using similar 200 μm thick longitudinal sections as above. Sections were stained with RedDot 2 nuclear stain (Biotium) and WGA Oregon Green (Fisher). At the Advanced Imaging Facility in the UC Davis School of Veterinary Medicine, a Leica TCS SP8 fit with a Mai Tai deep see laser was used for SHG microscopy. A 25x water immersion objective was used for imaging in conjunction with the multiphoton laser, tuned to 870nm and 830nm in series. Sections were imaged three times at random locations to obtain images stacks with a total and slice thickness of 100 μm and 1 μm, respectively.

A custom MATLAB script and processing in ImageJ was used to analyze the images stacks as previously described (14, 39)

### Trabecular and Cortical Bone Microstructural Properties

Following dissection, femurs were preserved in 70% ethanol and were imaged using micro-computed tomography (SCANCO Medical, µCT 35, Brüttisellen, Switzerland) to quantify trabecular bone microstructure at the distal metaphysis and cortical bone microstructure at the mid-diaphysis. Bones were embedded in 1.5% agarose during imaging and were scanned with a 6 µm isotropic nominal voxel size (x-ray tube potential = 55 kVp, current = 114 μA, integration time = 900 ms). Trabecular bone volumes of interest excluded the cortical shell and started adjacent to the metaphyseal growth plate extending 250 slices (1.5 mm) proximal. Trabecular bone volume fraction (BV/TV), trabecular thickness (Tb.Th), and other outcomes were determined using the manufacturer’s analysis software. Cortical bone volumes of interest included 100 slices (600 µm) centered at the longitudinal midpoint of each femur. Cortical bone area (B.Ar), total cross-sectional area (Tt.Ar), and other outcomes were determined using the manufacturer’s analysis software.

### Bone 3-point mechanical testing

Following µCT scanning, femurs were mechanically tested in three-point bending to determine structural and material properties of cortical bone. Bones were removed from 70% ethanol and rehydrated for 10-15 minutes in phosphate buffered saline (PBS) prior to mechanical testing using an electromagnetic materials testing system (ElectroForce 3200, TA Instruments, New Castle, DE) with the anterior aspect of each bone in tension. Data were collected at a sampling frequency of 50 Hz, the lower supports had a span of 5 mm, and the center loading plate was driven at 0.01 mm/sec until failure. Resulting force and displacement data were analyzed to determine structural properties of cortical bone (e.g., stiffness, yield force, ultimate force). Cross-sectional geometry at the mid-diaphysis determined from µCT was used to calculate material properties of cortical bone (e.g., elastic modulus, yield stress, ultimate stress) using standard beam theory equations.

### Statistical Analysis

Images were analyzed while blind to their genotype and treatment. Z-score analyses were done within each muscle and then combined into groups based on treatment (PBS or BAPN) or age (young or adult). Three-way ANOVAs in GraphPad Prism were run by genotype (wildtype or *mdx*), age (young or adult), and treatment (BAPN or PBS) for each muscle. In cases where data were separated by sex, three-way ANOVAs by genotype, age, and treatment were performed separately within male and female groups. Post-hoc Sidak multiple comparison tests were done in GraphPad Prism to discover differences between individual groups. Correlations and linear regressions were performed in GraphPad Prism on pairs of quantitative data. Stepwise multiple linear regressions were run in MATLAB with significance to enter the model set to p<0.10 and significance to exit the model set to p<0.20. Unless otherwise stated, significance was set at p<0.05.

## RESULTS

### Muscle mass and function

Prior to muscle isolation, the body weight of the mice was recorded. As expected, there was an increase in body mass in the adult mice compared to the young (Fig. 1A). BAPN did not have a consistent impact on body weight, but increased body weight in young *mdx* while leading to decreased body weight in adult *mdx* mice. All skeletal muscles of the *mdx* mice were impacted, but the diaphragm was grossly altered with highly fibrotic regions interspersed within muscle (Fig. 1B). We compared the limb muscle mass relative to the overall body weight and noted increases in both *mdx* soleus muscles compared to the wildtype soleus (Fig. 1C). As expected, the specific tension generated by the limb muscles was significantly reduced in both the soleus (Fig. 1D) and EDL (Fig. S1A) but was not affected by BAPN treatment in any muscle or age. The passive elastic muscle stiffness of the *mdx* diaphragm was significantly increased compared to wildtype as anticipated (Fig. 1E), but again BAPN had no impact on muscle stiffness. Neither did BAPN impact limb muscle elastic stiffness, although in soleus muscle there was a significant increase in elastic stiffness with age across conditions. The resistance to the initial stretch, dynamic stiffness, was similar to changes in elastic stiffness, but with an even more pronounced increase in the *mdx* diaphragm (Fig. S1B). The elastic index or ratio of elastic to dynamic stiffness thus decreased as *mdx* diaphragm became less elastic in mdx mice (Fig. 1F). The elastic index also decreased with age in *mdx* diaphragms. In the soleus however, the *mdx* muscles became significantly more elastic further indicating the muscle specific impacts. So, while the functional impact of *mdx* was muscle specific with decreased specific tension and increased stiffness in a muscle dependent manner as anticipated, treatment with BAPN did not induce significant changes to muscle function.

**Figure 1.**
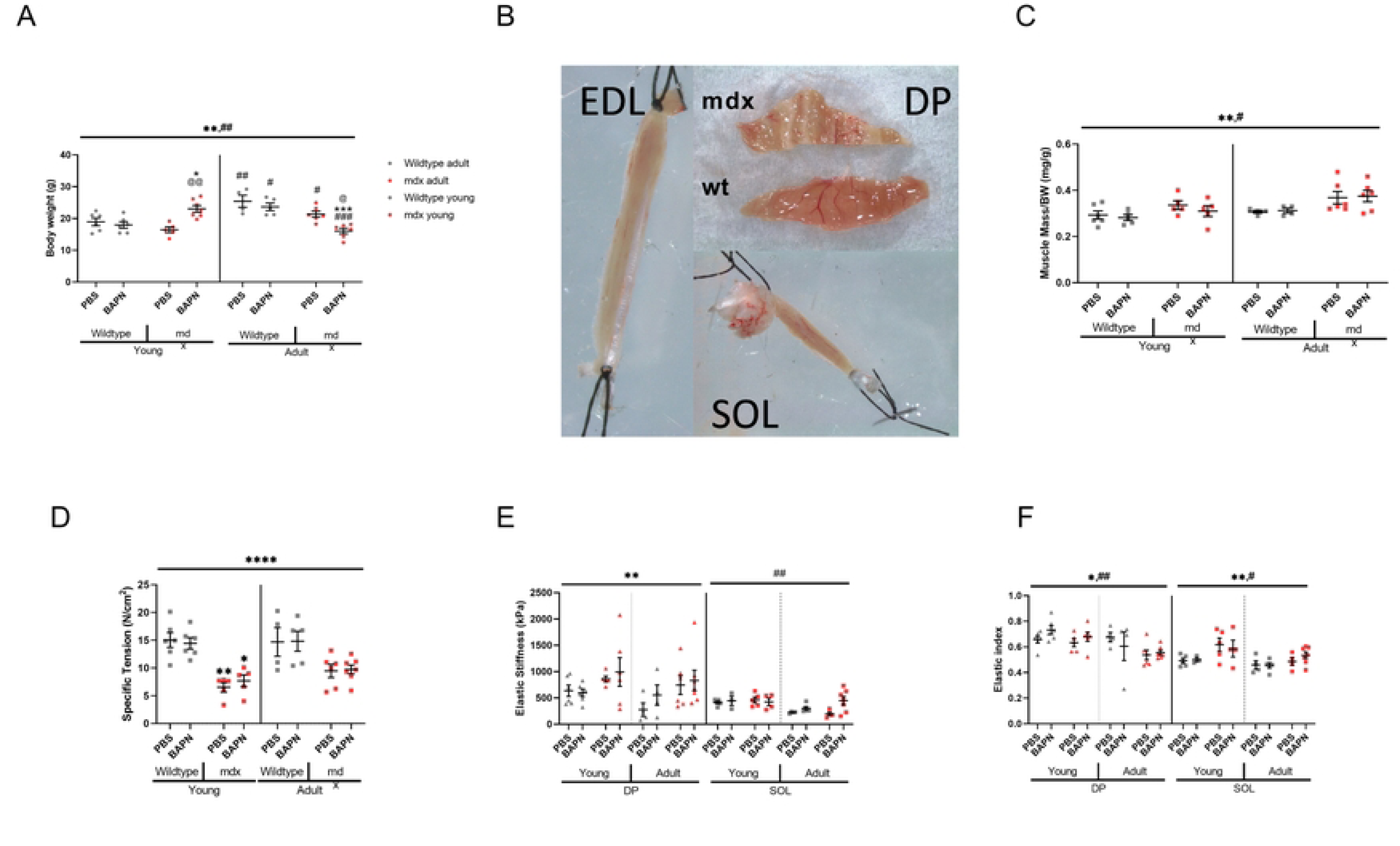
BAPN and Muscle Mechanics in wildtype and mdx. (A) Total body weight was greater for adult mice than young mice in both mdx and wildtype. (B) Representative image of isolated EDL, diaphragm (DP), and soleus (SOL) muscles for ex vivo mechanical testing. (C) In soleus, there were significant effects of age, genotype on muscle mass. (D) The specific tension of mdx soleus muscles were significantly lower than wildtype. (E) Elastic stiffness taken as the elastic modulus at 10% strain showed mdx DP was significantly stiffer than the wildtype diaphragms, and young soleus was significantly stiffer than adult soleus. (F) Elastic index, related to the amount of force remaining after stress relaxation compared with the peak force, showed significant effect of age and genotype in diaphragms, soleus. *=genotype effect, #=age effects, *=p<0.05, **=p<0.01, ***=p<0.001, ****=p<0.0001, determined by three-way ANOVAs with post-hoc Sidak multiple comparisons tests.

### Collagen content and cross-linking

As expected, there was a dramatic increase in collagen of the adult *mdx* diaphragm compared to wildtype with a significant, but moderate increase in collagen in the EDL (Fig. 2A). The wildtype diaphragm collagen didn’t increase into adulthood, but in the case of *mdx* there was a substantial increase as diaphragm fibrosis onset. In the EDL muscle, collagen increased with age for both genotypes. Insoluble collagen represents more maturely cross-linked collagen and followed a similar pattern of being increased in *mdx* and adult muscles and was significant in diaphragm and EDL muscles (Fig. 2B). The insoluble percentage is an index of the relative extent of collagen cross-linking. Collagen cross-linking was increased in the *mdx* mice diaphragms for both age groups and muscles. However, despite BAPN’s function to inhibit cross-linking, treatment did not impact relative collagen cross-linking for any group (Fig. 2C).

**Figure 2.**
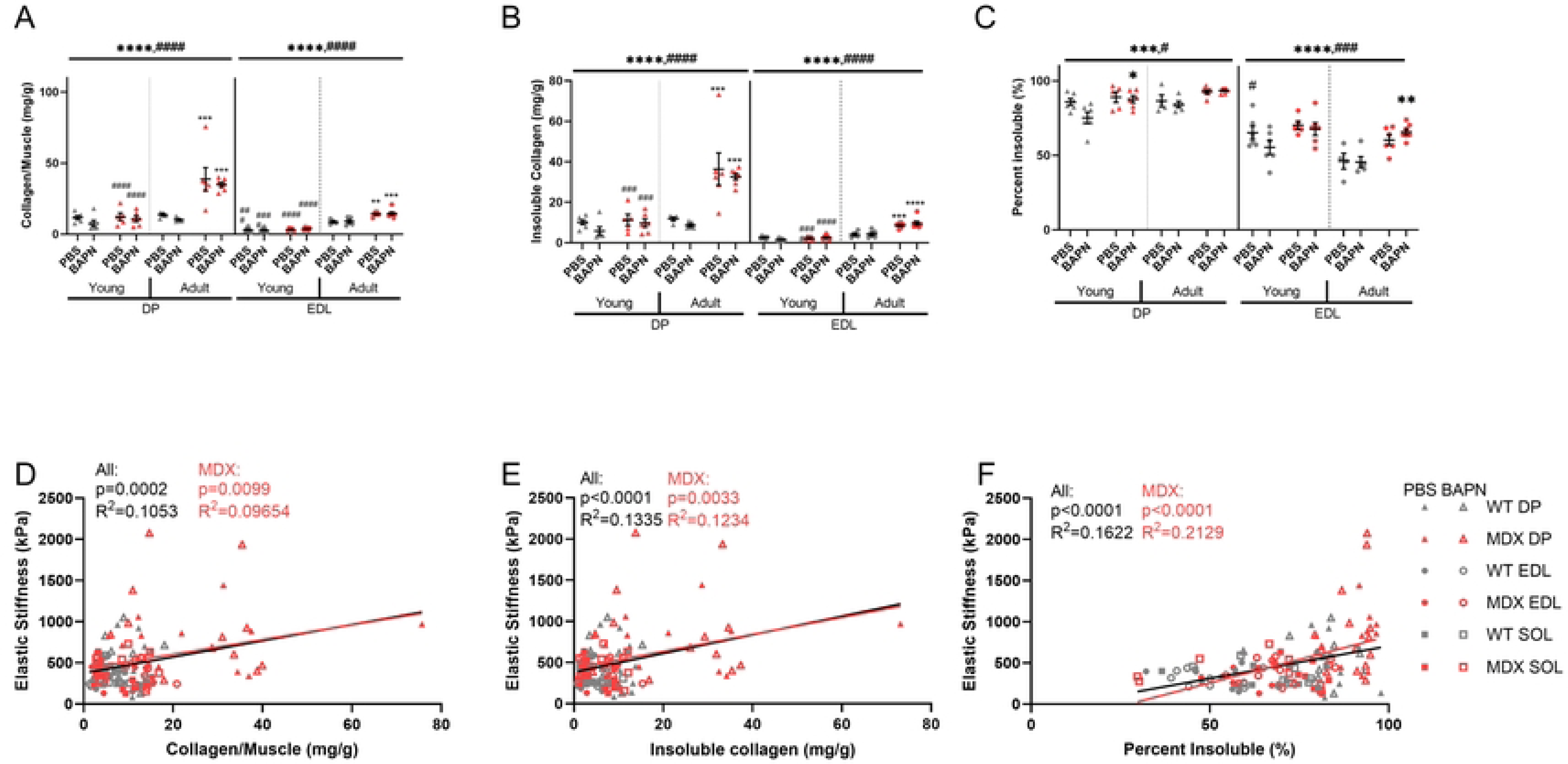
Collagen content and cross-linking. (A) The amount of collagen per muscle in wildtype and mdx muscles. The diaphragms and EDL muscle collagen content of mdx mice was significantly increased compared with the wildtype. (B) Insoluble (cross-linked) collagen was higher in mdx and adult muscles compared to wildtype. (C) The percentage of collagen that was cross-linked was higher in the mdx diaphragm and EDL muscles compared to the wildtype. Percent insoluble collagen was higher in the adult diaphragm and soleus and lower in the adult EDL compared to the respective young muscles. (D-F) Total collagen, insoluble collagen, and percent insoluble collagen scaled with elastic stiffness across all muscles together and the set of mdx points. *=genotype effect, #=age effects, *=p<0.05, **=p<0.01, ***=p<0.001, ****=p<0.0001, determined by three-way ANOVAs with post-hoc Sidak multiple comparisons tests.

Despite the ineffectiveness of BAPN in inhibiting collagen cross-linking in the muscle the natural variation in collagen and cross-linking could impact the measured elastic stiffness. Across muscles and within the *mdx* muscle in particular, there was a positive relationship between stiffness and total collagen (Fig. 2D), cross-linked insoluble collagen (Fig. 2E), and relative cross-linking as percent insoluble (Fig. 2F). The correlations were modest, but highly significant. Importantly, cross-linked collagen was more tightly associated than total collagen while relative extent of cross-linking was most well associated with stiffness. These associations support that collagen cross-linking contributes to muscle stiffness, but BAPN treatment was not effective in reducing collagen cross-linking.

### Collagen alignment with polarized light microscopy

Thick (200 μm) longitudinal vibratome cut sections were stained with Sirius red and imaged them with polarized light microscopy to look at alignment on a subpixel scale (MicroECM alignment) and across the section (MacroECM deviation) (Fig. 3A). There were no significant differences related to age, genotype, or BAPN treatment when looking at MacroECM deviation, the inverse of ECM alignment (Fig. 3B, Fig. S2B). Interestingly, there was a highly significant drop in MicroECM alignment in the adult diaphragms of both wildtype and *mdx* mice, but this was not observed in limb muscles (Fig. 3C, Fig. S2A). MacroECM deviation did not correlate with elastic stiffness overall (Fig. 3D) and neither did MicroECM alignment (Fig. 3E).

**Figure 3.**
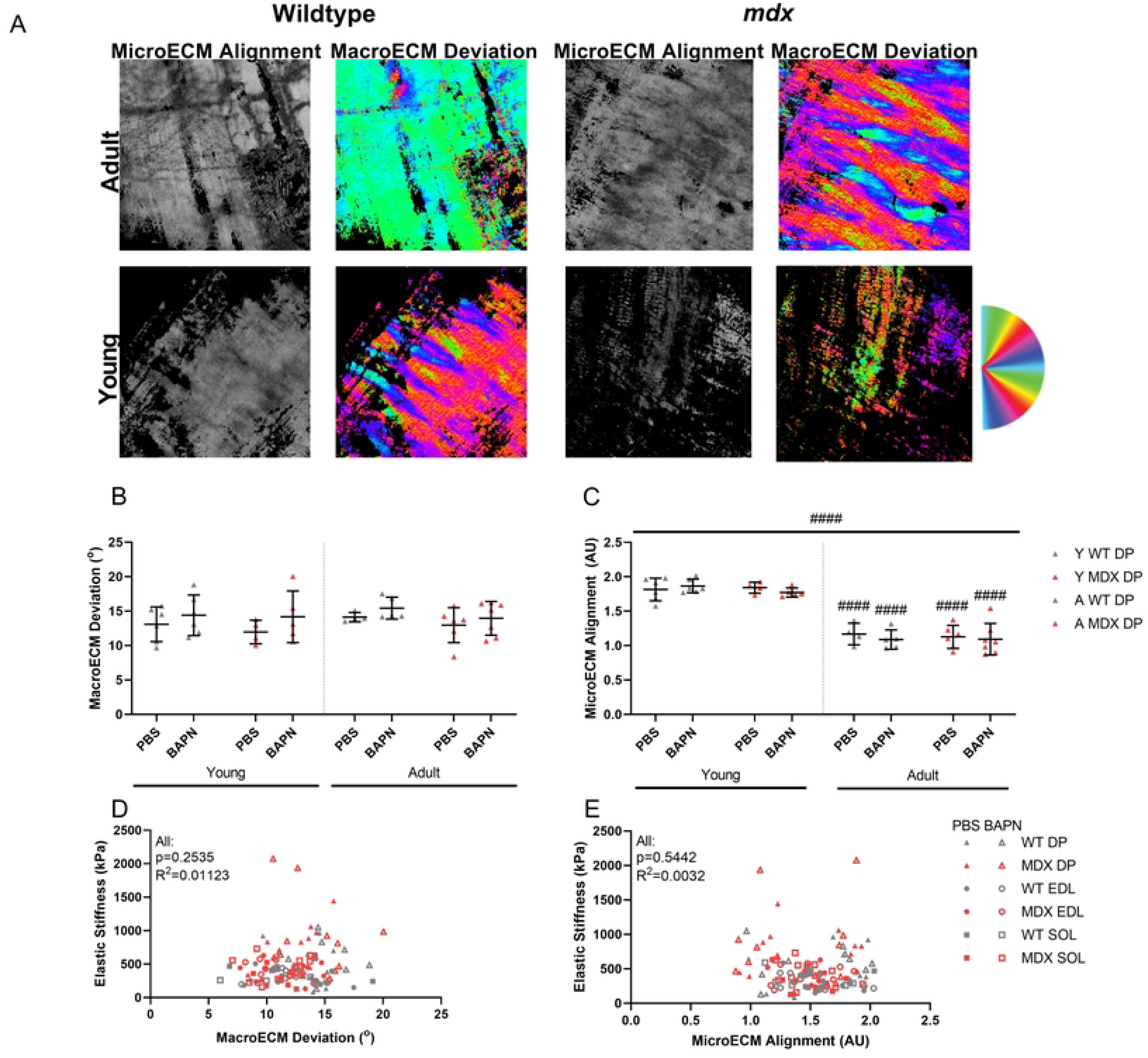
Sirius Red Collagen Architecture. (A) Representative images of MicroECM alignment (left) and MacroECM deviation (right) in diaphragm muscles. (B) The MacroECM deviation showed no significant differences between genotypes or ages in the diaphragm. (C) The quantification of the MicroECM alignment showed significant differences between young and adult in both wildtype and mdx diaphragms. (D) The relationship between MacroECM alignment and elastic stiffness showed no significant correlation overall or in individual muscles. (E) There was no significant relationship between MicroECM deviation and elastic stiffness overall or in any individual muscle. Significance bar represents significant main effect as determined by three-way ANOVA across treatment, genotype, and age. Significance symbol above groups indicates significance from corresponding genotype or age group according to post-hoc Sidak multiple comparisons tests. #=age effects, #=p<0.05, ##=p<0.01, ###=p<0.001, ####=p<0.0001.

Interestingly, MicroECM alignment did correlate with elastic index negatively in the soleus (R^2^=0.3140) (Fig. S2C). There was also a weak correlation between elastic index and MacroECM deviation across all muscles (R^2^=0.055) (Fig. S3D). Thus, polarized light microscopy did not identify any impacts of BAPN treatment, but demonstrated an age-related shift in diaphragm ECM microarchitecture.

### Collagen architecture with second harmonic generation microscopy

Representative images show different patterns of collagen orientation within the muscle with variable alignment, fiber size, and fiber area (Fig. 4A, Fig.S3A). Collagen deviation, the inverse of collagen alignment, was decreased in *mdx* EDL as previously observed (40) and adult soleus (Fig. 4B, Fig. S3C). However, the fibrotic diaphragm showed no differences in collagen deviation between age, genotype, or treatment. Collagen fiber area was significantly increased in the adult *mdx* diaphragm compared to the wildtype corresponding to the high collagen and severe fibrosis (Fig. 4C). While limb muscles collagen fiber area was not significantly changed by dystrophy, there were significant increases in collagen fiber area with age in the EDL and soleus (Fig. 4C, Fig. S3B). Collagen fiber size was not significantly different among groups in the diaphragm, but there was a significant effect of age such that there were larger collagen fibers in the adult soleus compared to the young (Fig. S3E). The collagen alignment could be a function of tissue strain, which can be captured by sarcomere length measurements obtained by SHG microscopy. There was a weak but significant positive correlation between sarcomere length and collagen deviation across all muscles and in the diaphragm (All: R^2^=0.0369, DP: R^2^=0.1210) (Fig. S3F), but with no changes in sarcomere length between groups the resting sarcomere length is unlikely to drive effects in this study (Fig. S3D). Collagen architecture measured by SHG was also significantly correlated with elastic stiffness, although the trends varied by muscle. Across muscles collagen deviation scaled positively with elastic stiffness (R^2^=0.0673), which was the opposite of the hypothesized relationship. Notably, in the EDL lower collagen deviation and thus more aligned collagen was correlated with higher stiffness as predicted (R^2^=0.1227) (Fig. 4D).

**Figure 4.**
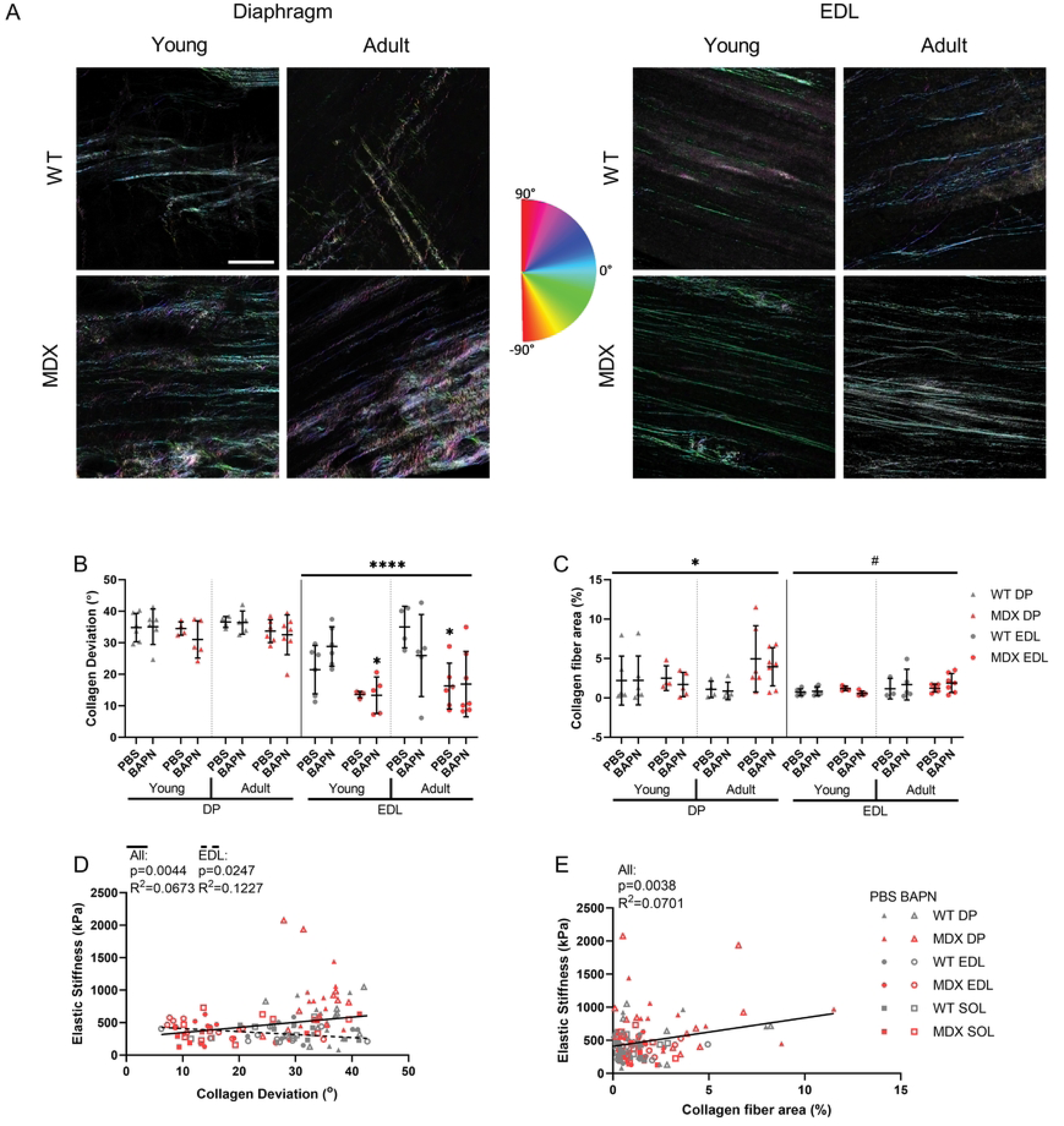
Collagen architecture is altered in dystrophic muscle and relates to passive mechanics. (A) Second Harmonic Generation (SHG) representative images of diaphragm and EDL muscle sections from young, adult, wildtype, and mdx mice. Colormap representsangles of each pixel window in the images as analyzed by OrientationJ. Scale bar is 100μm. (B) The mdx EDL had significantly lower collagen deviation (higher alignment) compared to the wildtype. (C) Collagen fiber area was increased in mdx diaphragms compared to wildtype. Adult EDL muscles had higher collagen fiber area than young EDL muscles. (D) All groups combined had a significant positive correlation of collagen deviation and elastic stiffness. The EDL had an independent significant negative correlation. (E) Collagen fiber area correlated positively with elastic stiffness across all muscles. Significance bars represent significant main effects determined by three-way ANOVAs for treatment, genotype, and age. Significance symbol above groups indicates significance from corresponding genotype according to post-hoc Sidak multiple comparisons tests. Significance by genotype: *p<0.05, **p<0.01, ***p<0.001, ****p<0.0001. Significance by age: #p<0.05.

On the other hand, collagen fiber area scaled positively with elastic stiffness overall (R^2^=0.0701) as would be expected, but the effect was not present within individual muscles (Fig. 4E). Direct visualization of collagen fibers again failed to show an impact of BAPN. Importantly however, this method did show unique adaptations by muscle where increased collagen alignment contributed to *mdx* EDL stiffness where more collagen fibers drive stiffness in adult *mdx* diaphragm.

### Relationships between collagen architecture and muscle function

To form a holistic view of how each ECM parameter related to functional parameters as well as other ECM parameters correlation matrices were made for all muscles and within each muscle group (Fig. S4A-D). While many significant relationships were observed, in order to determine which were most prominent and non-redundant multiple linear regressions were run across and within muscles. Over all muscles the most powerful predictor was the percentage of cross- linking while the amount of cross-linked collagen contributed independently as well (Fig. 5A). The collagen deviation entered into the model also, but surprisingly with more deviation, or less alignment, leading to more stiffness. These relationships can be driven by independent differences between muscles. Within the EDL in particular, the relationship to collagen deviation was as expected and previously observed with more alignment leading to more stiffness (Fig. 5B). The largest predictor in the EDL however, the area of collagen in perimysium fibers. In the EDL having less soluble non-cross-linked collagen was a predictor for stiffness likely related to less collagen cross-linking. In soleus muscles the relationship of elastic stiffness to collagen deviation was again reversed, albeit somewhat weaker (Fig. 5C). Notably, the diaphragm did not produce a significant model overall although in the adult diaphragm sarcomere length, collagen fiber area, and fiber size were all positive predictors of stiffness (Fig. S5E). Linear models were also run for adult and young muscles against elastic stiffness which showed fiber size, fiber area, and insoluble collagen as common positive predictors for stiffness (Fig. S5D-I). These models were comparable to those of dynamic stiffness which gave insoluble collagen and soluble collagen as common positive and negative predictors of stiffness, respectively (Fig. S5J-M). When linear models were made to predict specific tension in the EDL and soleus, the combined model gave insoluble collagen as a negative predictor of strength (Fig. S5A). The EDL model had collagen deviation as a contributor to specific tension in addition to less insoluble collagen (Fig. S5B) while the model for the soleus gave sarcomere length as a positive predictor for specific tension (Fig. S5C). Together, these relationships indicate that collagen architecture is more predictive of muscle stiffness than total collagen content with collagen cross-linking a consistent contributor while collagen alignment has a muscle specific influence on stiffness.

**Figure 5.**
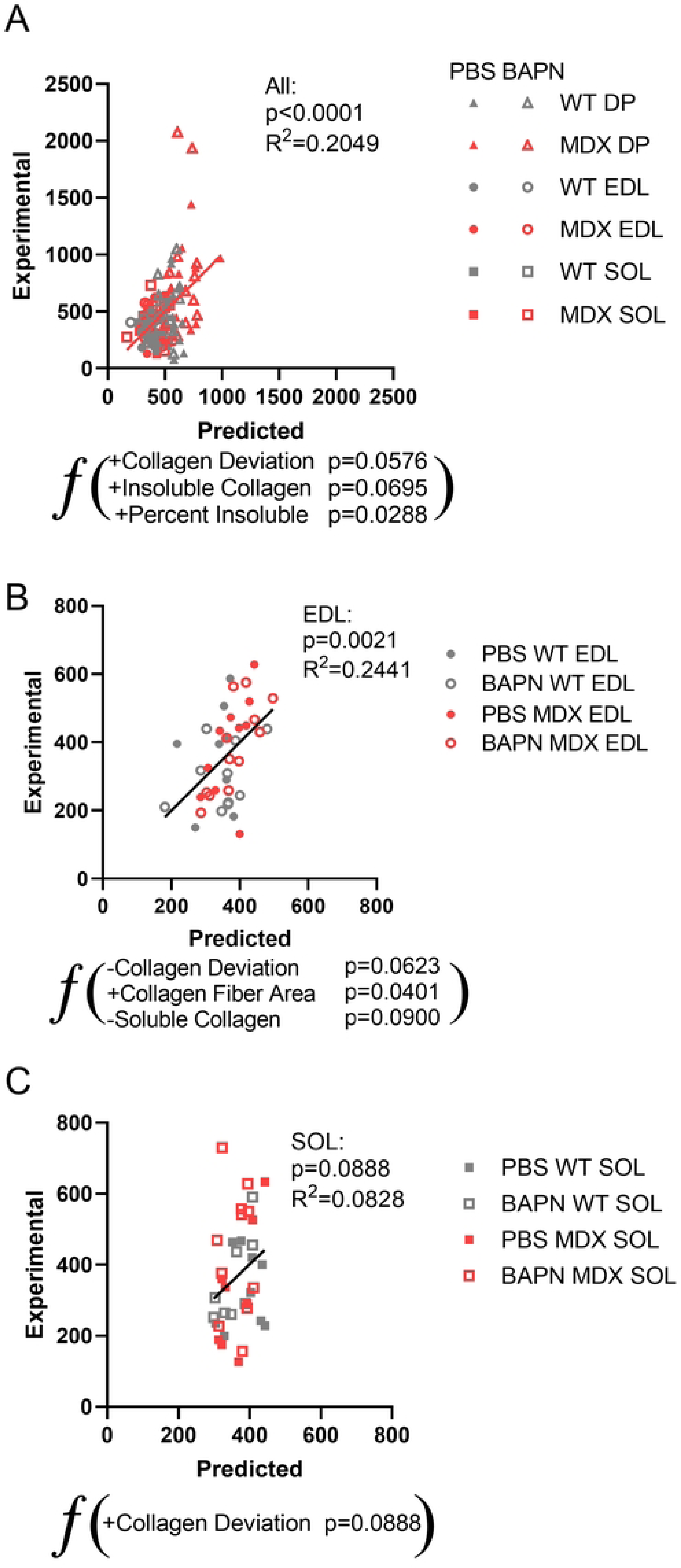
Multiple Linear Regressions show significant relationships between elastic stiffness and collagen architecture. (A) A multiple linear regression model was run in MATLAB to predict elastic stiffness across all muscles and individually within each muscle. The model produced significant predictors collagen deviation, insoluble collagen, and percent insoluble. (B) Collagen deviation and soluble collagen were negative predictors and collagen fiber area was a positive predictor of EDL elastic stiffness. (C) Collagen deviation was a positive predictor of elastic stiffness in the soleus.

### Dystrophic bone morphology and mechanics

Mid-diaphyseal total cross-sectional area of the femur determined by µCT (Fig 6A) showed substantial increase with age as expected, along with an increase in *mdx* mice compared to wildtype mice for both males and females (Fig 6B). However, BAPN did not have a large impact on bone cross-sectional area except potentially in the young *mdx* males. The cortical bone area had a similar effect of age, but was lower in *mdx* mice, particularly in adults (Fig 6C-D). Again, BAPN impact was isolated and in this instance to the young female *mdx* mice. Trabecular thickness also dramatically increased with age and was decreased broadly in *mdx* femurs (Fig. 6E). Importantly, BAPN significantly lowered trabecular thickness in female mice. Conversely, trabecular bone volume fraction did not change with age or BAPN treatment but was depressed in *mdx* mice (Fig. 6F).

**Figure 6.**
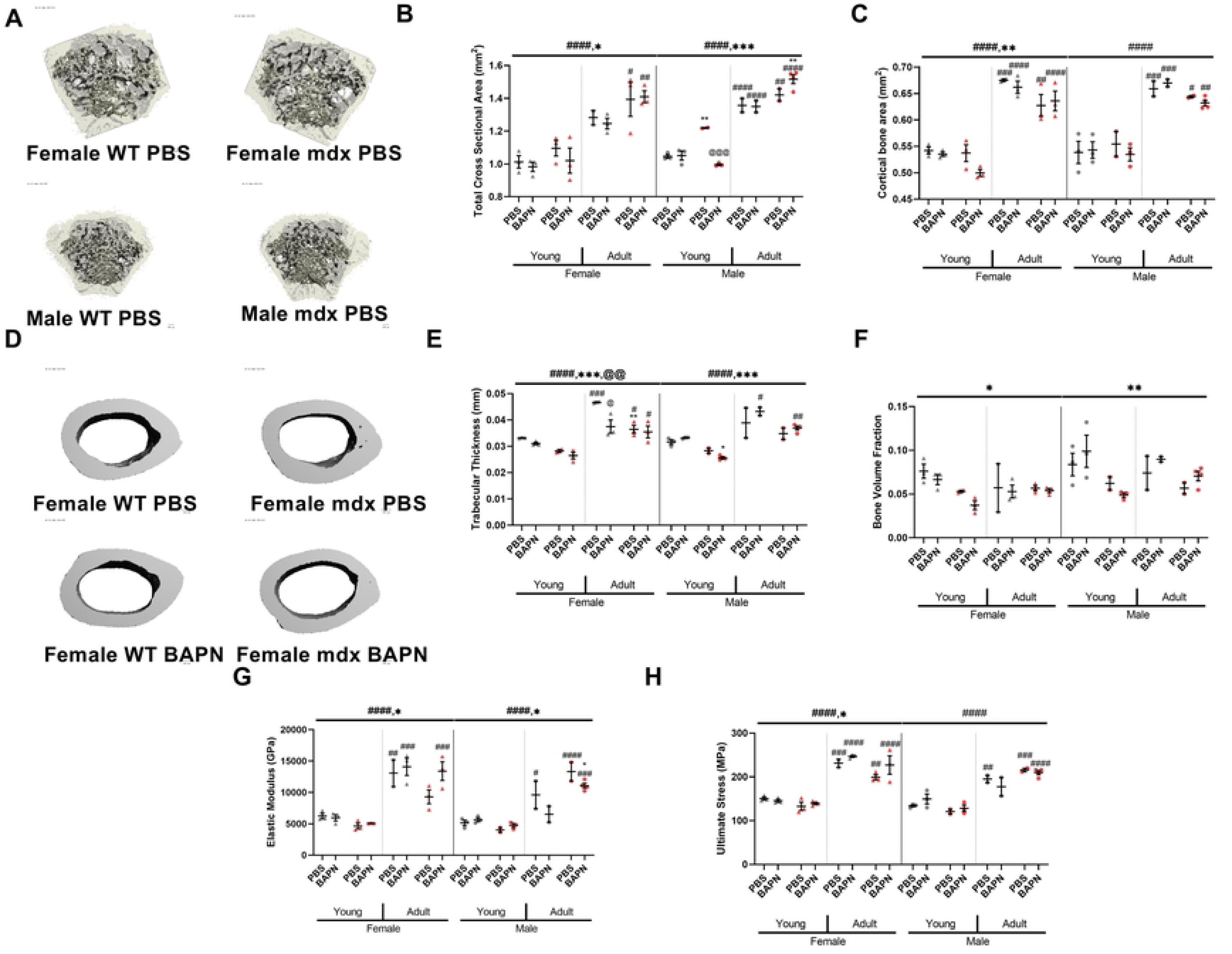
Bone. (A) Representative images of mid-diaphyseal total cross-sectional area of the femur by μCT (B) Total cross-sectional area was significantly increased in adult and mdx mice. (C) Cortical bone area was significantly increased in both male and female mice, and female mice also showed decreased area in mdx mice. (D) Representative images of cortical bone area. (E) There was an increase in trabecular thickness in adult mice, and a decrease in mdx mice. BAPN also decreased thickness in adult female wildtype mice. (F) Trabecular bone volume fraction was decreased in mdx mice. (G)Adult mice had significantly increased elastic modulus. (H) Adult mice had significantly increased ultimate stress. Significance by genotype: *p<0.05, **p<0.01, ***p<0.001, ****p<0.0001. Significance by age: #p<0.05, ##p<0.01, ###p<0.001, ####p<0.0001 Significance by treatment: @p<0.05, @@p<0.01, @@@p<0.001, @@@@p<0.0001 determined by three-way ANOVA test with post-hoc Sidak multiple comparisons tests.

Unsurprisingly, bones from adult mice exhibited substantially higher elastic modulus (Fig. 6G) and ultimate stress than in young mice (Fig. 6H). Female *mdx* mice had significantly lower stiffness and ultimate stresses, but in males the *mdx* effect was less consistent. Importantly, the modest impact of BAPN on bone structure did not contribute to any defect in bone mechanical properties. These bone analyses substantiate age related defects in *mdx* mice that are more prominent in females but with only a minor impact of BAPN on bone morphology with the applied treatment conditions.

## DISCUSSION

Although BAPN administration did not generate meaningful differences in skeletal muscle properties compared to PBS, there were notable shifts in the ECM architecture with dystrophy by muscle and age that have not previously been described. While expected differences were present with age such as greater body weight and muscle mass, there were notable distinctions such as lower elastic stiffness and elastic index in the adult muscles versus young. Two of the most robust age, genotype, and muscle-specific differences were the levels of collagen content and cross-linking. All muscles showed significant effects of age related to percent insoluble, insoluble collagen content, and total collagen while the EDL and diaphragm showed increases in those parameters in the *mdx* mice compared to wildtype. In all muscles the adult groups trended higher than the young for more cross-linked and total collagen content. This led to a higher proportion of collagen cross-linking in the adult diaphragm, but surprisingly, in the fast-twitch EDL the effect of age was reversed, with lower cross-linking in adult mice. Compared to limb muscles, the diaphragm showed the greatest amount of collagen content and cross-linking. This was greatly amplified in *mdx* diaphragm in line with previous studies showing severe fibrosis (14,16,41) and critical with diaphragm function dramatically compromised in DMD leading to progressive respiratory insufficiency (42, 43). Diaphragm muscles also show dramatic impairment in active mechanical properties not measured here (44, 45), but which are also compromised by fibrosis. This more robust characterization of the altered ECM architecture in the diaphragm emphasizes the need to discover anti-fibrotic therapies that effectively target the diaphragm in DMD to preserve diaphragm function throughout the lifespan.

The use of SHG microscopy was well suited to directly visualize the sarcomeric and collagen structure of the muscle. This study is the first to investigate the alignment of perimysial collagen fibers in the diaphragm. However, this aspect of architecture was not significantly altered in dystrophic mice as opposed to previous observation of limb muscles (14). Furthering the distinctions, across muscles increased alignment led to less stiffness. This was contrary to the previous investigation, but was driven by inclusion of the relatively high stiffness and high deviation of collagen in the diaphragm. The consistency of collagen alignment as a driving factor in the EDL stiffness further distinguished the uniqueness of the diaphragm. This points to the distinct relationship between collagen architecture and stiffness in different muscles. As with biochemical measures of collagen, the amount of perimysial collagen fibers measured by SHG were increased most prominently in the *mdx* diaphragm, but not until adulthood and not in the limb muscles. There is evidence that the collagen cables in the perimysium are important to the overall muscle mechanical properties and undergo changes in fibrosis that confer increases in tissue stiffness (14,46–48). While the amount of collagen in skeletal muscle is not always a consistent predictor of tissue stiffness (14,15,49), collagen fiber area seems to be relevant to the mechanical properties in some muscles given its incorporation in the multiple linear regressions of the overall EDL, adult EDL, and adult diaphragm and as demonstrated previously (14).

Collagen fiber size was a positive predictor for elastic stiffness in multiple linear models including the adult overall, adult diaphragm, and young EDL groups. While perimysial collagen fibers are only a subset of overall collagen, the importance of collagen fiber area and the alignment of those collagen fibers in determining stiffness highlight their important role in passive muscle function and as targets for anti-fibrotic therapies.

A wrinkle to the mechanism of dystrophic damage of the diaphragm includes its fan shape that incurs strains in multiple directions. One possibility is that given its low level of collagen alignment compared to the limb muscles, the diaphragm’s stiffness may be more dependent on its level of collagen content and cross-linking. In the multiple linear regression for elastic stiffness, the adult diaphragm included sarcomere length, collagen fiber area, and collagen fiber size as positive predictors while the regression for dynamic stiffness generated insoluble collagen as the only positive predictor for the overall and adult diaphragm. These models show ECM parameters of collagen cross-linking, fiber area, and fiber size as useful predictors for diaphragm stiffness, although the relevance of these parameters across aging is still unclear. Paired with the observation that the diaphragm collagen fibers are less aligned than those of the EDL and soleus, it may be the case that this alignment or lack thereof may play a role in the diaphragm’s stiffness in a different axis. This could also help explain why diaphragm alignment does not significantly relate to stiffness since we mechanically tested the diaphragm muscle along the myofiber angle. While some experiments have been done to estimate or directly measure diaphragm mechanics in the transverse axis (50, 51), further experiments testing diaphragm mechanical properties in multiple directions of strain and relating them to collagen architecture would further elucidate this relationship.

This study was designed to block collagen cross-linking in skeletal muscle. BAPN has been previously shown to be an effective collagen cross-linking inhibitor in multiple tissues (23, 52) and similar effects in muscle were anticipated. The dosage of 100 mg/kg/day and intraperitoneal delivery method were similar to previous experiments showing effectiveness of BAPN in blocking collagen cross-linking (53, 54). The moderate impact on bone morphology and trabecular thickness in particular supports that the BAPN dosage was active to some extent.

However, studies looking at the impact of BAPN in bone have used higher concentrations, such as 200 mg/kg/day or 500 mg/kg/day (24). Thus, increasing the dose may be necessary to elicit collagen cross-linking inhibition in musculoskeletal tissues (55, 56). Injecting the BAPN locally into the muscle may more effectively inhibit collagen cross-linking by allowing for high local concentrations, but repeated injections could be a significant source of injury. Alternative collagen cross-linking inhibitors may be more effective in skeletal muscle. The usage of neutralizing antibodies to the collagen cross-linking enzyme, lysyl oxidase like 2, have been shown to block collagen cross-linking in lung, liver, and kidney fibrosis (57–61). While BAPN was ineffective at blocking collagen cross-linking in this study, the results still advocate for effective inhibition of collagen cross-linking as an efficacious way to reduce muscle stiffness and contractures.

There are several aspects of our study that limit the interpretations of the data. As with all mouse models of DMD the severity and degree of fibrosis is less than the human condition. However, we utilized a more severely fibrotic mouse model with the *mdx* mutation on the DB2 background to produce a more fibrotic phenotype (26, 62). Another limitation is the linear strain placed on the diaphragm compared to how the diaphragm functions *in vivo*. Bi-axial investigation of the diaphragm passive mechanics may yield new insights into the unique collagen architecture. While measuring collagen architecture and mechanics in the same muscles provides greater power, the imaging is only performed on a small section of the muscle. For polarized light imaging few differences were observed, and the change in micro-alignment with age specifically in the diaphragm is difficult to interpret. For SHG-based measures, only the perimysium collagen fibers were analyzed, while contributions from the epimysium and endomysium were not incorporated. Despite these limitations this study provides the most comprehensive investigation of skeletal muscle passive mechanics dependence on ECM architectural features to date.

Despite the inability to evaluate the influence of reduced collagen cross-linking on muscle mechanics as intended, this study provides a robust analysis of ECM features and muscle stiffness across dystrophy, age, and muscles. This led to discovery that while the EDL muscle stiffness is closely related to collagen alignment is a major factor in muscle stiffness, dystrophic diaphragm muscle stiffness is more dependent on increased collagen content and cross-linking. This emphasizes that while the ECM response to muscular dystrophy is relatively consistent across muscles, with more collagen, cross-linking, and alignment, the implications for stiffness are muscle dependent. This study furthers our knowledge of the determinants of passive muscle stiffness and is critical for future efforts evaluating anti-fibrotic therapies in models of muscular dystrophy for their impact on muscle function and stiffness.

## ACKNOWLEDGEMENTS

This work was supported by a grant from the NIH National Institute of Arthritis and Musculoskeletal and Skin Diseases (NIAMS; R00AR067867). The UC Davis Health Sciences District Advanced Imaging Facility and Dr. Ingrid Brust-Mascher supported the SHG Imaging. The work was supported by the lab Dr. Keith Baar including access to equipment and attentive comments on the study. We would like to thank Xinyue Li for support in preparation of the manuscript and thoughtful discussions. We would also like to thank additional members of the MyoMatrix Lab at UC Davis including CJ Mileti, Taryn Loomis, Daryl Dinh, and Matthew Nakaki for insightful conversations and review of the data.

## DATA AVAILABILITY

A spreadsheet with all the annotated data for this manuscript is available from figshare under the DOI (10.6084/m9.figshare.20234712)

## SUPPLEMENTAL FIGURE LEGENDS

Figure S1. Muscle tetanus in EDL and Dynamic Stiffness. (A) Muscle specific tension was significantly decreased in mdx EDL muscles. (B) Dynamic stiffness was increased in mdx in the diaphragms and was also increased in young in the soleus compared to adult. *=genotype effect, #=age effects, @=treatment effects, *=p<0.05, **=p<0.01, ***=p<0.001, ****=p<0.0001, determined by two-way ANOVAs with post-hoc Sidak multiple comparisons tests.

Figure S2. Sirius Red collagen architecture relates to muscle elasticity. (A-B) There were no significant main effects determined by three-way ANOVAs across treatment, genotype, and age for EDL and soleus muscle MicroECM alignment and MacroECM deviation. (C) MicroECM alignment of overall and mdx soleus muscles scaled negatively with elastic index. (D) MacroECM deviation demonstrated a weak positive correlation with elastic index across all muscles.

Figure S3. (A) Second Harmonic Generation (SHG) representative images of soleus muscle sections from young, adult, wildtype, and mdx mice. Colormap represents angles of each pixel window in the images as analyzed by OrientationJ. Scale bar is 100μm. (B) Adult solei had higher collagen fiber area than young solei. (C) Adult and mdx solei had lower collagen deviation (higher alignment) than young and wildtype solei. (D) Adult diaphragm muscles had longer sarcomere lengths than young diaphragms, and wildtype solei had longer sarcomere lengths than mdx. (E) Adult soleus muscles had larger mean collagen fiber diameters than young soleus muscles. (F) Collagen deviation had a positive correlation with sarcomere length across all muscles and in the diaphragm. (G) There was a significant positive correlation between sarcomere length and elastic stiffness across all groups. (H) There was no significant correlation between collagen fiber diameter and elastic stiffness. Significance bars represent significant main effects determined by three-way ANOVAs for treatment, genotype, and age. Significance symbol above groups indicates significance from corresponding genotype or age according to post-hoc Sidak multiple comparisons tests. Significance by genotype: *p<0.05, **p<0.01, ***p<0.01; significance by age: #p<0.05, ##p<0.01.

Figure S4. Correlation matrices demonstrated significant linear relationships between end-point parameters across and within muscles. (A) Correlation matrix showing significant relationships between parameters across all muscles. (B-D) Correlation matrices of individual muscles showing significant relationships between parameters in diaphragm, EDL, and soleus muscles. *p<0.05, determined by Pearson correlation analysis.

Figure S5. Multiple Linear Regressions show significant relationships between collagen architecture and specific tension/passive stiffness. (A) The combined model produced insoluble collagen as a negative predictor of specific tension. (B) The model gave collagen deviation as a positive predictor for specific tension in EDL and insoluble collagen as a negative predictor. (C) The model gave sarcomere length as the only positive predictor for specific tension in soleus. (D-I) Multiple linear regressions of young and adult groups of muscles demonstrated significant predictors of elastic stiffness. (J-M) Multiple linear regressions of combined and individual muscles showed collagen architecture relates to dynamic stiffness.

## Notes

### Competing Interest Statement

The authors have declared no competing interest.

